# A spectral demixing method for high-precision multi-color localization microscopy

**DOI:** 10.1101/2021.12.23.473862

**Authors:** Leonid Andronov, Rachel Genthial, Didier Hentsch, Bruno P. Klaholz

## Abstract

Single molecule localization microscopy (SMLM) with a dichroic image splitter can provide invaluable multi-color information regarding colocalization of individual molecules, but it often suffers from technical limitations. So far, demixing algorithms give suboptimal results in terms of localization precision and correction of chromatic aberrations. Here we present an image splitter based multi-color SMLM method (splitSMLM) that offers much improved localization precision & drift correction, compensation of chromatic aberrations, and optimized performance of fluorophores in a specific buffer to equalize their reactivation rates for simultaneous imaging. A novel spectral demixing algorithm, SplitViSu, fully preserves localization precision with essentially no data loss and corrects chromatic aberrations at the nanometer scale. Multi-color performance is further improved by using optimized fluorophore and filter combinations. Applied to three-color imaging of the nuclear pore complex (NPC), this method provides a refined positioning of the individual NPC proteins and reveals that Pom121 clusters act as NPC deposition loci, hence illustrating strength and general applicability of the method.

## Introduction

Super-resolution microscopy breaks the diffraction limit of resolution in fluorescence microscopy and hence allows bioimaging with unprecedented details. Among the numerous super-resolution imaging techniques, stimulated emission depletion (STED) and structured illumination microscopy (SIM) rely on special illumination schemes^1,2^. Single-molecule localization microscopy (SMLM) techniques include stochastic optical reconstruction microscopy (STORM), photoactivated localization microscopy (PALM) and points accumulation for imaging in nanoscale topography (PAINT). They use rather conventional epifluorescence schemes and rely on specific photophysical and photochemical properties of dyes that are repeatedly switched on and off, imaging only a sparse subset of molecules at a time^3–5^. MINFLUX and similar techniques combine both approaches and localize individually switched-on fluorophores with a patterned illumination^6–9^. SMLM is one of the most heavily used bioimaging super-resolution techniques because it employs conventional optics and simple labelling^10^ and yet provides excellent resolution in the 10-20 nm range, usually higher than what is achievable with STED and much higher than with SIM, hence allowing imaging at the molecular level.

An important advantage of fluorescence imaging is its multi-color capability. This allows for colocalization of objects through simultaneous imaging of different targets in a given sample. While classical colocalization in confocal microscopy is limited in resolution, super-resolution microscopy combined with multi-color imaging can help deciphering interactions between proteins at the single molecule level^11–13^. Multi-color imaging is possible in SIM and STED using fluorophores with substantially different spectral properties^14^ or lifetimes^15^. SMLM, however, opens additional possibilities for multi-color imaging, thanks to the access to the fluorescence properties of individual molecules, enabling separation of fluorophores with very close spectra using a dichroic image splitter and ratiometry^16^ or single-molecule spectrometry^17^. Potential advantages include: 1) several species of labels are acquired simultaneously, improving the imaging speed; 2) thanks to simultaneous imaging, drift is the same in all channels and can be corrected more reliably; 3) spectrally close fluorophores often have similar photophysical properties and can provide equivalent data quality for different color channels; 4) non reliable localizations originating from autofluorescence or noise can be filtered out; 5) chromatic aberrations can be avoided. However, while the potential of multi-color SMLM is strong, few implementations actually allow to do this conveniently and precisely.

Separating fluorophores with similar spectra can be achieved with a dichroic mirror that divides the imaging path of the microscope into two channels, the short wavelength (λ_S_) channel and the long wavelength (λ_L_) channel, followed by simultaneous recording of the two channels on the camera^18–20^ (**Fig. 1**, **Suppl. Fig. S1**). This can be implemented with minor modifications of a microscope, *e.g.* using a commercial image splitter such as Optosplit (Cairn Research) or W-View Gemini (Hamamatsu Photonics), where the two channels are imaged side-by-side on the same camera chip (**Fig. 1C**). For each fluorophore, the ratio of its photon counts in the two channels is *r=I_L_/I_S_*. It will depend on its emission spectra and transmission (reflection) spectrum of the dichroic mirror, providing different ratios for fluorophores even with only slightly different emission spectra (**Fig. 1D-E**).

**Figure 1.**
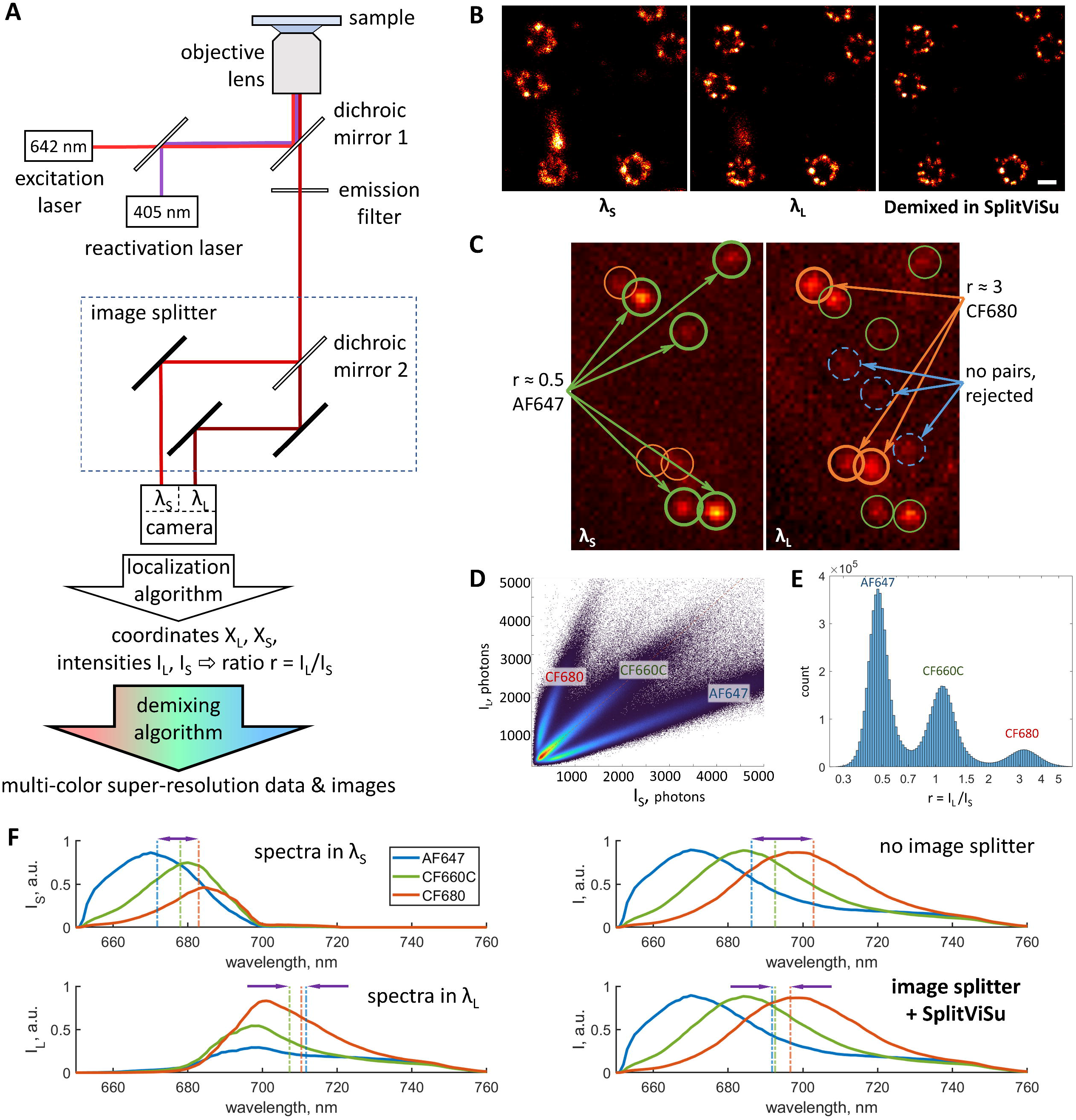
Principle of splitSMLM. (**A**) Scheme of the splitSMLM microscope. (**B**) SR images of a single-labelled sample, reconstructed from localizations in the λ_S_ channel, λ_L_ channel or after demixing in SplitViSu, demonstrating removal of spurious localizations and resolution improvement. (**C**) Image of the two spectrally different channels, λ_S_ & λ_L_, with localizations originating either from AF647 (green), CF680 (orange) or from background noise (blue). (**D**) Bivariate histogram of photon counts *I_S_* & *I_L_* originating from the λ_S_ & λ_L_ channels allowing separation of three fluorophore species. (**E**) Univariate histogram of ratios *r* on a semi-log plot with peaks corresponding to different fluorophore species. (**F**) Fluorescence spectra of AF647, CF660C and CF680 observed within the λ_S_ & λ_L_ channels, without image splitter, or equivalent spectra after demixing in SplitViSu. Vertical lines represent average detected wavelengths for the corresponding fluorophores in the λ_S_ & λ_L_ channels, respectively, without image splitter, or after demixing in SplitViSu. Arrows show maximum spectral difference within the triplet of fluorophores and reflect the amount of chromatic aberrations in the corresponding setting. Scale bar, 100 nm.

The coordinates and the brightness of the fluorophores within the split camera image can be determined using a conventional localization algorithm^21^, while additional processing is needed to assign the localizations to the fluorophores. There are a few tools available for demixing^22,23^ that implement a rather simple algorithm to identify localization pairs from two spectral channels, followed by fluorophore assignment based on their intensities within the channels, and listing of demixed coordinates using the coordinates of one of the input channels.

The precision of single-molecule localization increases with the square root of the number of detected photons^24^. In order to achieve the highest resolution, it is therefore important to detect and keep as many photons as possible. At the same time, the dichroic mirror in the image splitter strongly increases spectral shift between fluorophores detected in different channels (**Fig. 1F**), which increases problems with chromatic aberrations. For this reason, using an image splitter, the coordinates of unmixed localizations are commonly calculated from only one spectral channel^25,26^. However, this approach leaves the photons in the second channel unused and therefore reduces the localization precision. The photons in the second channel are then used only for spectral demixing by the ratio of photons between two channels. Because the signal of the same channel is used for image reconstruction in different colors, this approach reduces chromatic aberrations (**Fig. 1F**, **Suppl. Fig. S2**). To improve the localization precision, it is possible to widen the spectral width of the channel used for localization and to narrow that of the second channel^9,26^. This, however, decreases the signal within the narrow-band channel, which affects the reliability of demixing of spectrally close fluorophores and leads to a high cross-talk or to an increased rejection of localizations, especially with strong background typical for bioimaging. New approaches for spectral demixing taking into account drift and chromatic aberration corrections are therefore needed.

Among the best fluorophores for SMLM so far are the molecules based on the cyanine scaffold with emission in the far-red and near-infrared spectral ranges of 620-750 nm^27^. These include Alexa Fluor 647, CF680, CF660C, DY643, DY654, DyLight 680^26–28^. Their close excitation and emission spectra do not allow conventional multi-color imaging where each dye is excited with a separate laser and is detected through a separate emission filter. However, they are perfectly suited for excitation with a single laser and for simultaneous imaging into two channels through a dichroic splitter (**Fig. 1F**, **Suppl. Fig. S1B**). Similarly to the photophysics of other fluorophores, the emission spectra of these dyes have a rapid increase that starts near their excitation maximum, followed by a longer tail after the emission peak (**Suppl. Fig. S1B**). In order to efficiently discriminate their spectra, it is important to detect the spectral region where they differ most, *i.e.* the region between their excitation and emission maxima. At the same time, this region lies closely to the excitation laser line and is usually rejected by emission filters. This not only rejects precious emitted photons but also makes the spectral discrimination less reliable. Detection of emission within this spectral region (“salvaged fluorescence”) has been shown to be beneficial for spectral separation but up to now required a rather complex optical setup^26^.

Finally, sample preparation is a key factor because imaging buffers play a huge role in the performance of fluorophores in SMLM^29–31^ with rather variable performances for different fluorophores^30–32^. Since in multi-color SMLM the performance of fluorophores affects not only their localization but also their spectral separation, it is important to choose buffers to maximize the photon yield of the fluorophores and yet obtain comparable efficiencies for each fluorophore.

In this work, we describe a simple and robust implementation of multi-color SMLM with a spectral image splitter (splitSMLM) resulting in much improved localization precision, drift correction, compensation of chromatic aberrations and performance of fluorophores in optimized buffers. To implements the spectral demixing data processing for multi-color SMLM we developed a software, SplitViSu, which is freely available and is integrated into a workflow of image analysis, reconstruction and segmentation of SMLM data within the SharpViSu platform^33^. We demonstrate the performance of our methods by imaging the nuclear pore complex (NPC), a reference standard for super-resolution microscopy^34^, revealing simultaneous localization of individual proteins within the nuclear envelope and providing new findings on their colocalization.

## Results

Among the various far-red fluorophores suitable for multi-color SMLM, we chose three fluorophores: Alexa Fluor 647 (AF647), CF660C and CF680^35^, because they have a similar cyanine-based structure, similar but sufficiently different spectra and are widely available as antibody conjugates. Additionally, they have already been characterized and performed well for multi-color SMLM in the classical Glucose Oxidase - Catalase buffer with addition of β-mercaptoethylamine (MEA) or β-mercaptoethanol.

For spectral demixing, we use a bivariate histogram of intensities in the λ_L_ and λ_S_ channels (**Fig. 1D**, **Suppl. Fig. S3A-C**). On such a histogram, the localizations originating from a given fluorophore are distributed along a line with slope *r* and zero y-intercept: *I_L_ = r* · *I_S_*, or if represented on a log-log graph (**Suppl. Fig. S3C**), along a line with slope *1* and y-intercept *log(r)*: *log(I_L_) = log(r) + log(I_S_)*. The bivariate histogram has an advantage over a univariate histogram of ratios *r* (**Fig. 1E**, **Suppl. Fig. S3D**) because it allows demixing or filtration based not only on ratios but also on absolute values of intensities. Because spectral demixing becomes less reliable at low intensities, a bivariate histogram allows for removal of such low-intensity localizations that cannot be reliably assigned to a fluorophore.

### Maximizing the localization precision and minimizing chromatic aberrations in multi-color imaging

The coordinates of the demixed localizations are typically calculated using input coordinated of only one spectral channel^16,25^, thus losing the photons detected in the second channel. In splitSMLM, we aimed at improving the localization precision by including the signals from both channels into the computation of the coordinates of the unmixed fluorophores.

Let us regard each single-molecule localization as a normally distributed random variable X = N (μ, σ^2^), where mean μ is the position of the fluorophore and the standard deviation σ = σ_0_/√I, where σ_0_ is the standard deviation of the point spread function and I is the number of detected photons. With a dichroic image splitter, for each molecule, there are two experimental localizations, X_L_ = N(μ_L_, σ_L_^2^) and X_S_ = N(μ_S_, σ_S_^2^), that originate from the the λ_S_ and the λ_L_ channels, respectively. In an ideal image splitter without losses, all input photons are split into two channels with a dichroic mirror: I = I_L_ + I_S_. The ratio r=I_L_/I_S_ depends on the spectral characteristics of the mirror and on the emission spectra of the fluorophore. The two positions of a fluorophore imaged through the splitter do not coincide due to chromatic aberrations: μ_S_ ≠ μ_L_ (**Fig. 1F**, **2A**, **Suppl. Fig. S2**). We will neglect here the chromatic aberrations between fluorophores acquired through the same channel, because the average wavelength is very close for different fluorophores within one channel (**Fig. 1F**, **Suppl. Fig. S2**). The demixing algorithm decides, based on I_L_ and I_S_, to which species the localization belongs, and then calculates the position of the unmixed localization based on its X_L_ and X_S_.

**Figure 2.**
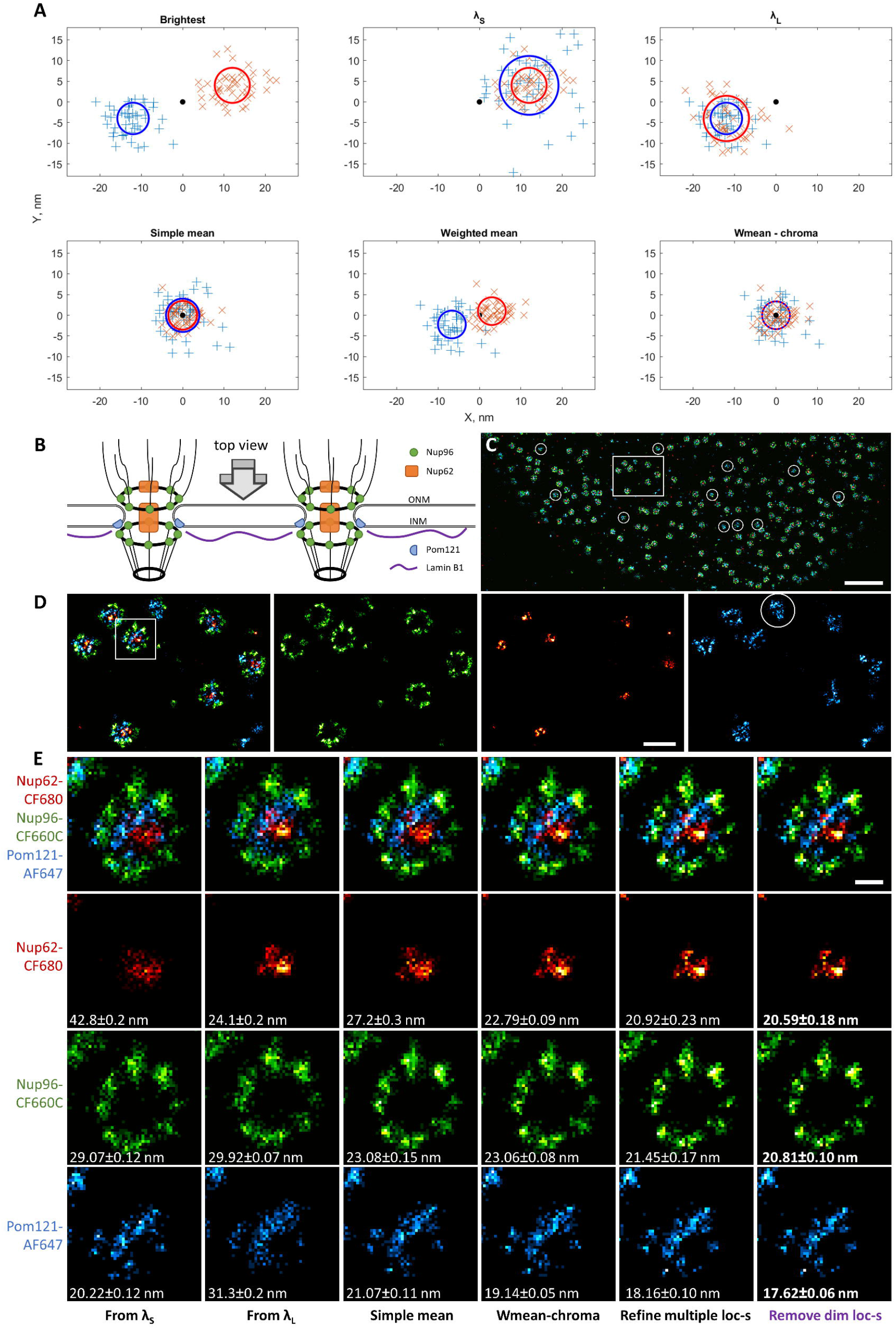
Localization precision and chromatic aberration correction in splitSMLM using SplitViSu. (A) Simulated results of demixing in SplitViSu using various methods for calculation of the output coordinates: “brightest” – the output coordinates equal the input coordinates from the brightest channel for the given fluorophore; “λ_S_”, “λ_L_” – the output coordinates equal the input coordinates from the λ_S_ or λ_L_ channel, correspondingly; “simple mean” – the output is calculated as a simple mean of the input coordinates in the λ_S_ or λ_L_ channels; “weighted mean” – the output is calculated as an photon count-weighted mean of the input coordinates in the λ_S_ or λ_L_ channels; “wmean - chroma” – the output is calculated as the “weighted mean” with subsequent subtraction of chromatic aberrations. The simulated data contains two species of fluorophores with different ratios: r_red_ = 0.6, r_blue_ = 3.5; the true position is the same for both fluorophore species and is shown as a black dot; the total photon number detected from each localization is the same for all fluorophores and equals 2000 photons; the full width at half maximum (FWHM) of the microscope PSF equals to 300 nm; the chromatic aberrations between two fluorophores within the opposite channels of the image splitter equal to 24 nm (x-direction) and 8 nm (y-direction); each species was detected 50 times at both channels. The circles represent FWHM of the demixed distributions with corresponding settings. (B) Scheme of the NPC’s at the NE, used as a test structure in this study. ONM, outer nuclear membrane; INM, inner nuclear membrane. (**C-E**) “Top views” of the NPC’s in a U2OS cell with immunofluorescently labelled Pom121 (blue), Nup62 (red) and Nup96 (green). Rectangle in C represents the region zoomed in in D; square in D represents the region zoomed in in E. Circles in C & E represent Pom121 clusters with few localizations of Nup96 and Nup62. (**E**) Different methods for calculation of demixed coordinates tested on a single NPC. Numbers in the bottom represent the resolution of the images, calculated according to the FRC_1/7_ criterion^36^. The images in (E) are reconstructed as bivariate histograms of localization coordinates with a histogram bin size of 5×5 nm. Scale bars, 1 µm (C), 200 nm (D), 50 nm (E).

In the case where all unmixed localizations adopt coordinates from the λ_L_ channel, X = X_L_ = N(μ_L_, σ_L_^2^), there are no chromatic aberrations (μ = μ_L_), but the localization precision is decreased: σ = σ_0_/√I_L_ = σ_0_/√(I – I_S_) > σ_0_/√I. A similar result is obtained when all localizations adopt coordinates from the λ_S_ channel (**Fig. 2A**). To have best localization precision (less than √2 times worse than when all photons are used for localization), one would choose for localization the channel that has higher intensity. In many cases, however, the channel with highest intensity for one fluorophore will be the channel with lowest intensity for another fluorophore. Now, if one chooses to localize this second fluorophore within the channel that has highest intensity, the chromatic aberrations will come into play because different species will be localized from spectrally different channels. One would often rather “sacrifice” the localization precision in order to avoid chromatic aberrations and would lose more than √2 times in localization precision for the fluorophores that have less intensity in the channel used for localization^16,25^.

In the following, we describe a way to utilize the photons present in both channels for both spectral demixing and localization. We developed an algorithm to calculate the coordinates of the unmixed localization as a weighted sum of its coordinates in the two spectral channels:

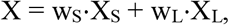

where w_S_ and w_L_are intensity-dependent weights that can adopt values from 0 to 1:

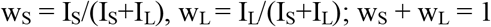

The resulting localizations are normally distributed X = N(w_L_⋅μ_L_ + w_S_⋅μ_S_, w_L_^2^⋅σ_L_^2^ + w_S_^2^⋅σ_S_^2^) and are localized in between of the two initial localizations, thus resulting in reduced chromatic aberrations (μ = w_L_ μ_L_ + w_S_⋅μ_S_). The chromaticity is not completely removed though, because the position of the unmixed molecules becomes dependent on the ratio of their intensities in the λ_S_ and λ_L_ channels:

X = X_S_/(1+r) + r ⋅ X_L_/(1+r), where r = I_L_/I_S_. It is corrected better for fluorophores that have closer *r*-values (fluorophores with closer emission spectra).

Importantly, the localization precision in this case equals that for imaging without image splitter:

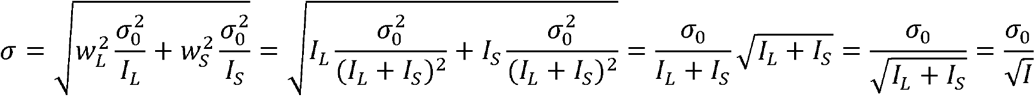

Therefore, the weighted mean method fully preserves the localization precision and yet reduces chromatic aberrations (**Fig. 2A**).

Now let us consider a case where the unmixed coordinates are calculated as a simple mean of X_S_ and X_L_: X = (X_S_ + X_L_) / 2 = N((μ_L_ + μ_S_)/2, (σ_L_^2^ + σ_S_^2^)/4). The chromatic aberration would be completely corrected for all localizations – the unmixed localization is located in the middle between the two input localizations: μ = (μ_L_ + μ_S_)/2. The localization precision, however, would be decreased, unless I_L_ = I_S_:

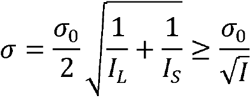

In practice, it is important to know under which conditions the localization precision using the ‘mean’ method will be higher than when using the coordinates of only the brightest channel. Suppose I_L_ > I_S_, then the condition for the improvement of the localization precision can be written as:

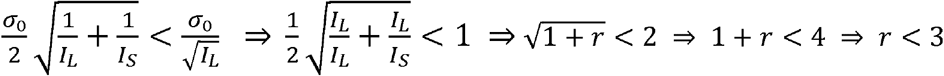

The localization precision using the ‘mean’ coordinates will be improved if the ratio of the intensity between the λ_S_ and λ_L_ channels for the given fluorophore is less than 3, which is the vast majority of cases in practice. It should be noted that using this method, the localization precision is always improved by a factor > √2 if a fluorophore has most of its emission in the channel that is not used for localization but only used for demixing in the previously reported demixing algorithms.

The coordinates calculated by the “weighted mean” method exhibit residual chromatic aberrations equal to the chromatic aberration between the fluorophores imaged without image splitter (**Suppl. Fig. S2**), plus an unequal magnification of the channels that might be produced by the image splitting optics. For a given fluorophore with a constant ratio *r*, the residual shift depends only on the fluorophore’s position within the image and can therefore be registered using information already available in the data. We propose to register the shift with respect to the position, calculated using the “simple mean” method, as it provides the position closest to the ground truth for all fluorophores (**Fig. 2A**, **Suppl. Fig. S2**). For each fluorophore within a dataset, we calculate its simple and weighted mean positions, and their difference: Δx = x_wm_-x_m_. Δx has a random component due to the limited localization precision and due to fluctuations in the photon counts, and a component that depends on the position x due to the chromatic aberrations: Δx = Δx_rand_ + Δx_chro_. The typical values for our setup and fluorophore triplet are: - 20 nm ≤ Δx_rand_ ≤ 20 nm, −4 nm ≤ Δx_chro_ ≤ 4 nm (**Suppl. Fig. S4**). We fit the chromatic component Δx_chro_ as a function of the coordinates with a polynomial, separately for x and y coordinates, and subtract it from the “weighted mean” positions: X = x_wm_ -Δx_chro_ (**Suppl. Fig. S4**). The resulting localizations have the best possible resolution, as determined by Fourier ring correlation^36^ (**Fig. 2E**) and, importantly, no noticeable chromatic shift (**Fig. 2**).

### Refinement of multiple localizations for SMLM image reconstruction

For SMLM image reconstruction, we developed a new method for processing of relocalizations of molecules on consecutive frames. Usually, consecutive localizations are merged into a single value that is located at the mean position of the initial localizations. This improves its precision but reduces the density of localizations that might not necessarily result in resolution improvement (**Suppl. Fig. S5**). In our method, instead of being reduced to only one localization, all initial localizations are kept, but their coordinates are refined. The new coordinates are selected randomly from a normal distribution with a standard deviation σ = σ_psf_/√N_ph_, where σ_psf_ is the standard deviation of the point spread function of the microscope (σ_psf_ = 140 nm in our setup) and N_ph_ is the sum of the photon counts of the series of consecutive localizations. This is done to reflect new (*i.e.* improved) localization precision and to avoid appearance of very bright pixels in the reconstructed images, which would happen if all localizations of the group were set to have exactly the same coordinates^21^. The center of the refined distribution lies at the weighted mean position of the initial localizations, the weights of the initial localizations being proportional to their photon counts. This method improves the resolution of the reconstructed images, both visually and according to the FRC criterion (**Fig. 2E**). After the refinement, the localizations with low photon numbers can be additionally removed, which further improves resolution (**Fig. 2E**). This method of processing of consecutive localizations is now implemented in the software suite SharpViSu^33^.

### SplitViSu, a software for demixing of spectrally close fluorophores

To integrate different aspects of fluorophore colocalization analysis into a convenient tool, we developed SplitViSu, a software for demixing SMLM data acquired with an image splitter. The software can be used as a stand-alone application for Windows, or can be run under Matlab or from SharpViSu^33^. It allows for visualization of input data, flexible selection of the regions for each channel, automatic or manual alignment of localizations between the channels, pairing of localizations within a given distance, visualization of the paired localizations as a bivariate histogram of intensities, flexible assignment of the localizations to fluorophores based on the photon counts and their ratios, and estimation of cross-talk between unmixed localizations.

### Spectral properties of filters for image splitting optics

In this work, we managed to detect the “salvaged fluorescence” region by carefully selecting filters on conventional epifluorescence setup that allowed to detect fluorescence starting from 652 nm, only 10 nm apart from the excitation laser and to better discriminate fluorophores with very close emission spectra (**Suppl. Fig. S7**). To separate the emission light into two channels, we used a dichroic mirror at 690 nm, that provides, for the used fluorophores, similar intensities and background in both channels (**Fig. 1F**, **Suppl. Fig. S1B**), allowing to use a single run of a conventional fitting algorithm for both channels. For detection spectrally close to a laser line, conventional filters usually do not provide low enough background for single-molecule observation. Each combination of filters must be carefully designed and tested. In our setup, we therefore used an additional band-pass filter in the λ_S_ channel of the image splitter in order to remove residual background from the laser (**Suppl. Fig. S1A**).

In a setup where the emission spectra are split into a narrow λ_S_ band that is used for spectral discrimination and a broad λ_L_ channel that receives most of the fluorescence and is used for localization^26^, the fluorophore brightness differs a lot between the channels. This would lead to an under-detection of fluorophores in the λ_S_ channel, especially the species with longer-wavelength emission spectra. For fluorophores with emission in the shorter wavelength range, a substantial number of photons will still be lost in the λ_S_ channel. In our setup, we use λ_S_ and λ_L_ channels of similar spectral width that provides similar intensities of fluorescence and background in both λ_S_ and λ_L_, which allows performing localization with the same parameters for both channels. Moreover, in our processing algorithm all available photons are used for both localization and demixing.

### Removal of unreliable localizations by spectral detection

Interestingly, detection of individual fluorophores through a spectral image splitter is beneficial even for single-labelled samples (**Fig. 1B**). Firstly, because a given fluorophore species has a unique ratio *r=I_L_/I_S_*, the localizations with a different ratio might originate from a different species of dyes (*e.g.* from fluorophores that cause autofluorescence) or from spurious localizations (**Suppl. Fig. S6C-D**). Secondly, in our setup, we introduced a condition that in order to be considered for the analysis, a molecule is detected in both channels independently. A small tolerance (50-100 nm) between the channels is necessary to account for small chromatic aberrations and the localization imprecision in the input data. This provides a strong filter to reduce artefactual localizations originating from background noise or from poorly blinking regions (**Fig. 1B**, **Suppl. Fig. S6**). The localizations originating from background noise can be detected at random places, therefore they are unlikely to be detected at the same place in both channels at the same time, which provides a way to exclude them. Poorly blinking regions, depending on the localization algorithm, can cause erroneous localizations at different positions within two channels, they might not fall within the tolerance and would be removed. Also, multiple blinking involving more than one fluorophore species usually has a different *r* value and can also be filtered out.

It should be noted, however, that for samples with three and more fluorophores the removal of unreliable localizations becomes less efficient because the “unreliable” region for one fluorophore may coincide with the “reliable” region for the other dye.

### Buffer for simultaneous multi-color imaging

We also optimized the buffer composition because it can influence SMLM quite significantly. While the classical Glox-MEA buffer provides already acceptable performances for image splitter-based SMLM^28^, the presence of slowly-blinking low-intensity localizations of CF680 and CF660C^28^ is problematic because molecules cannot be reliably unmixed and should be rejected. Moreover, we find that AF647, CF660C and CF680 in this buffer demonstrate different reactivation responses on illumination with 405 nm light: AF647 can be reactivated continuously with gradual increase of the reactivation intensity (*e.g.*, in the 0 to 50 mW range, see methods), while CF660C and especially CF680 are reactivated well with very low power of the laser, but cannot be reactivated anymore when the power increases to intermediate or high values (**Suppl. Fig. S8B**). Because the reactivation power usually increases gradually during an acquisition, this effect leads to a non-constant ratio between the dyes during the experiment, *i.e.* there are more CF680 molecules at the beginning and more AF647 localizations at the end of the acquisition. This might lead to undesirable effects, for example, when the drift is not perfectly corrected or when, to save time, the acquisition has been stopped before complete bleaching of all three fluorophores.

We tested a variety of conditions and found that addition of 2 mM of cyclooctatetraene (COT) to the Glox buffer increases the photon yield of all three fluorophores by 20-50% without decreasing the density of localizations (**Suppl. Fig. S8A**, **S8C**), an effect also observed for AF647^29^. Moreover, we show that COT equalizes the response of these fluorophores on reactivating light. In Glox with COT, AF647 slightly decreases its response at high 405 nm intensities, while CF660C and CF680 substantially increase it (**Suppl. Fig. S8B-C**). COT increases the proportion of bright molecules for all three fluorophores (**Suppl. Fig. S8**) and allows for more reliable demixing and reduces rejection of localizations (**Suppl. Fig. S9**).

### Application of optimized three-color imaging on the nuclear pore complex

We applied our method for simultaneous three-color imaging of individual proteins within the nuclear envelope (NE) of human cells. Thanks to the ellipsoidal shape of the NE, one can observe the NPC’s projected under different angles: “top views” when focusing on the NE close to the coverslip and “side views” when focusing on the middle of the nucleus, several micrometers away from the coverslip (see scheme, **Fig. 2B, 3A**). NPC’s have dimensions of ∼150 nm and a partially symmetrical structure that makes it a perfect test object for super-resolution microscopy^34^. Within the NPC, a number of different proteins can be detected, providing a range of distances that can be used as nanometer-scale rulers for various imaging and labelling tests. We chose to label NPC subunits corresponding to distinct parts of NPC (see **Fig. 2B, 3A**) as well as lamin B1, a protein found close to the inner nuclear membrane.

**Figure 3.**
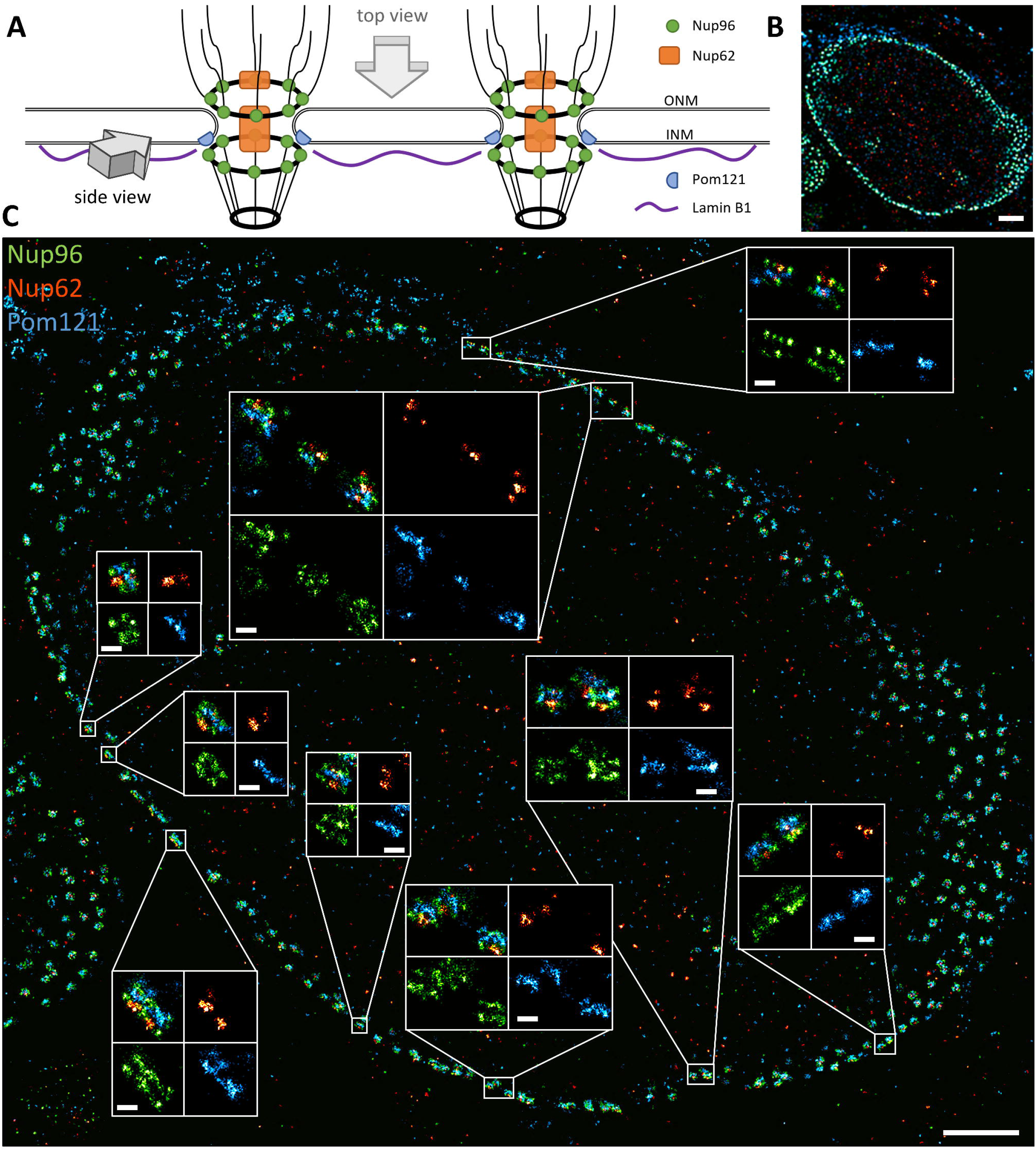
“Side view” of the NPC’s with Nup96, Nup62 and Pom121, imaged by splitSMLM. (**A**) Scheme of the NPC’s at the NE. ONM, outer nuclear membrane; INM, inner nuclear membrane. (**B**) Overview of a U2OS cell with immunofluorescently labelled Pom121 (blue), Nup62 (red) and Nup96 (green). (**C**) The cell from (B) with zoomed in “side view” regions in insets. Scale bars, 2 µm (B), 1 µm (C), 100 nm (insets).

Our SMLM images of Nup96 demonstrate 8-fold rotational symmetry with respect to the axis of the NPC and two-fold symmetric localization with respect to the NE (**Fig. 2-4**), in line with the structure of the NPC (**Fig. 2B**) and previous super-resolution studies^34^. Nup62 appears in the center of the NPC in the “top view” of our super-resolution images (**Fig. 2, 4A**), as can be expected for a central channel protein of the NPC. In the “side view”, however, it can be seen at both the nucleoplasmic (weaker signal) and the cytoplasmic sides (stronger signal) of the complex (**Fig. 3C, 4B**), indicating the presence of its copies at both the nuclear and the cytoplasmic sides of the NPC.

**Figure 4.**
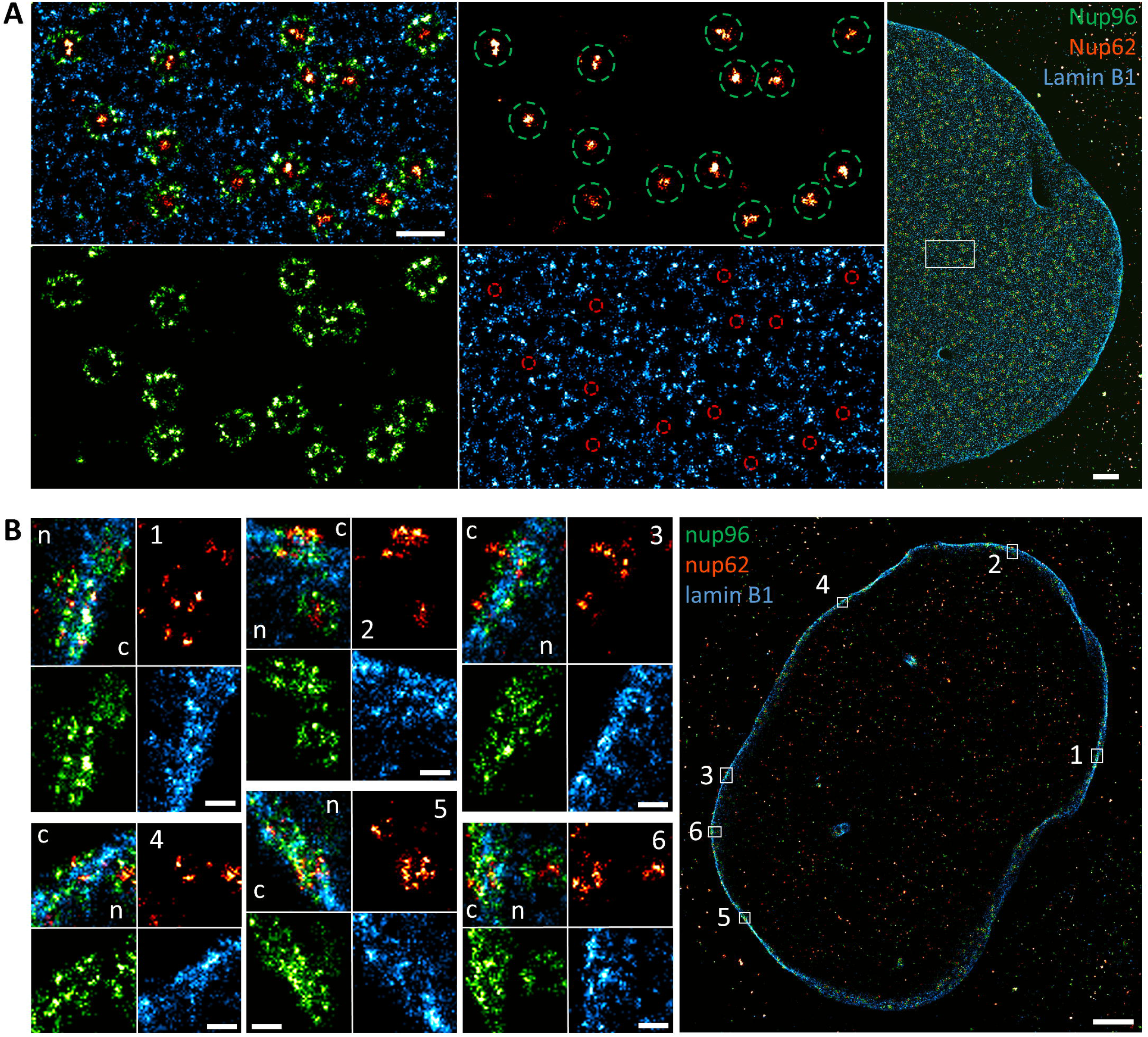
Three-color splitSMLM imaging of the nuclear pore complex. (**A**) “Top view” of the NE (Nup96, green; Nup62, red; lamin B1, blue). Green circles have a diameter of 150 nm and represent the positions of the Nup96 ring. Red circles have a diameter of 50 nm and represent the positions of Nup62. White rectangle represents the zoomed in region. (**B**) “Side views” of the NE. The numbered rectangles in the general view represent the zoomed in regions; n, nucleus; c, cytoplasm. Scale bars, 200 nm (A, left side), 1 µm (A, right side & B, right side), 100 nm (B, left side).

Our data suggests that Pom121 is located only at the nuclear side of the NE, close to the nuclear ring of Nup96 (**Fig. 3C**), confirming the role of Pom121 as an anchoring point for the NPC at the inner nuclear membrane^37,38^. In the “top view”, Pom121 forms an irregular ring structure between the regular Nup96 ring and the NPC central channel, revealed by the Nup62 staining (**Fig.2D-E**). The Pom121 localization inside the NPC in this projection corresponds to the deep protrusion of the NE into the NPC between the cytoplasmic and the nuclear rings of the Y-complexes^39^ (**Fig. 2B**). From the “top view” of our three-color images it is evident that lamin B1 is abundant at the NE, but it is completely absent precisely where NPC’s are located (**Fig. 4A**), which is in line with the current model of the inner nuclear membrane region (**Fig. 3A**). In the “side view”, however, we find that lamin B1 can be found in between the nuclear and the cytoplasmic rings of the NPC (**Fig. 4B**), closer to the cytoplasm than thought according to current models^40^.

## Discussion

Simultaneous multi-color SMLM with spectral image splitting was demonstrated in the early days of super-resolution microscopy^18,19^, but it did not spread much beyond experimental setups. In this work, we describe splitSMLM, an image splitting method for multi-color SMLM using a spectral image splitter, a dedicated software for spectral demixing (SplitViSu), a selected fluorophore triplet with 3 close wavelengths and an optimized buffer system that keeps all 3 fluorophores active in an equivalent manner. Thanks to simultaneous imaging and improved signal-to-noise ratios as essentially all photons are used, splitSMLM makes drift correction much easier, which is crucial for precise colocalization studies. The splitSMLM implementation is simple and robust and results in much improved localization precision, drift correction, compensation of chromatic aberrations at the nanometer scale and performance of fluorophores in optimized buffers.

The setup is built with commercially available components and is therefore easy to implement at biology laboratories. Previous implementations had to make compromises in terms of photon budget, using only a subset of detectable light for localization and therefore losing precision. Moreover, the low brightness and sub-optimal photo-switching of previously employed fluorophores led to under-detection and strong rejection of localization events, resulting in labelling density loss. The splitSMLM method presented here overcomes many of these limitations because it allows simultaneous rather than consecutive multi-color SMLM acquisitions without the need of separate acquisitions for each fluorophore at longer and shorter excitation and emission wavelengths, respectively^13,41,42^. Because high-quality SMLM requires long acquisitions (minutes to hours)^28^, the new method significantly reduces the acquisition time as compared to sequential imaging. The fact that the drift is the same in all channels allows summing all channels for drift estimation. Moreover, the present approach uses a specifically combined spectrally distinct fluorophore triplet (AF647, CF660C, CF680) for which conjugated antibodies are available and which achieves equally high performances thanks to an optimized buffer that is identical for the entire experiment. The proposed imaging with three fluorophores addresses the majority of cases needed in practice.

An important point to consider in the splitSMLM method is the combination of optimized choice of the fluorophore triplet and optimally adapted emission filters, which allows detecting fluorescence very close to the laser line (close to 10 nm), thereby achieving “salvaged fluorescence” with maximized detection and improved spectral separation of the fluorophores. The dichroic image-splitting mirror was chosen to split the emission into two channels with similar spectral widths, allowing reliable detection of molecules within both channels simultaneously. Furthermore, the optimized imaging buffer that comprises a combination of COT and Glox buffers increases the photon yield of all three fluorophores by 20-50% without decreasing the density of localizations (**Suppl. Fig. S8A**, **S8C**). This effect was previously demonstrated only for AF647^29^. COT turns out to equalize the response on reactivating light, thus allowing colocalization analysis under rather stable imaging conditions. Because COT increases the proportion of bright molecules for all three fluorophores, it is strongly beneficial for multi-color SMLM imaging as it improves demixing and reduces rejection of localizations (**Suppl. Figs. S8** & **S9**). The optimized imaging buffer composition hence improves the photon budget and the amount of reactivation of far-red fluorophores, which is beneficial for splitSMLM.

To analyse splitSMLM data conveniently, we developed a new demixing software, SplitViSu. This tool allows visualization of input data, flexible selection of the regions for each channel, automatic or manual alignment of localizations between the channels, pairing of localizations within a given distance, visualization of the paired localizations as a bivariate histogram of intensities, flexible assignment of the localizations to fluorophores based on the photon counts and their ratios, estimation of (unwanted) cross-talk between unmixed localizations (**Suppl. Fig. S10**). The software can be used as a stand-alone application under Windows, or can be run under Matlab or from SharpViSu^33^ in which it was added as a plugin. The demixing algorithm uses all detectable photons both for demixing and localization, improves the localization precision and corrects chromatic aberrations. The usage of similar wavelengths has the specific advantage here of reducing chromatic aberration thanks to the capability of the software to demix spectrally close fluorophores, which is improved even further by additional processing in SplitViSu. Moreover, we introduced the refinement of re-localization events, which further increases image resolution. Additionally, we show the benefits of an image splitting setup and fluorophore demixing also for single-labelled samples (**Fig. 1B, Suppl. Fig. S6**) thanks to filtering out localizations that originate from background noise or from autofluorescence. Unlike other software with some similar functionalities^22,23^, the SplitViSu tool provides flexibility in calculation of output localizations, with the possibility of freely selecting options to use only coordinates from a selected channel or use their mean or weighted mean values, thus improving the localization precision and compensating chromatic aberrations. Moreover, SplitViSu is fully integrated with SharpViSu^33^, allowing drift correction, visualization, estimation of resolution, filtration, cluster analysis with ClusterViSu and 3DClusterViSu^12,43^ and further processing of spectrally demixed multi-color data.

The strength of the splitSMLM method described here relies on a strong synergistic effect of several parameter that is obtained from the much-improved multi-color super-resolution data quality, achieved by refining each aspect of multi-color imaging - optics, data processing and performance of fluorophores. The quality improvements include 1) higher image resolution; 2) higher density of localization; 3) absence of cross-talk between the labels; 4) absence of chromatic shifts; 5) suppression of localizations originating from background noise or from autofluorescence.

We applied our new approach to three-color super-resolution imaging of the NPC and the nuclear lamina. The obtained splitSMLM data validate our experimental approach, confirming as expected^34^ the typical 8-fold localization of Nup96 in the “top view” and its symmetrical localization with respect to the NE in the “side view” (**Fig. 2-4**). Furthermore, the central localization of Nup62 within the NPC (**Fig. 2-4**) and absence of lamin B1 at the nuclear pores in the “top view” (**Fig. 4A**) now achieve a refined positioning of the individual proteins within the nuclear membrane as compared to earlier data^44^. Three-color imaging with Pom121 has not been done previously (only with single-color microscopy^45,46^) and now reveals the presence of irregular ring structures between the Nup96 ring and the central channel, represented by Nup62 (**Fig. 2D-E**). In the “side view”, it is visible that Pom121 localizes only at the nuclear side of the NPC (**Fig. 3D**), which suggests that Pom121 clusters act as NPC deposition loci at the inner nuclear membrane, providing unprecedented insights not seen in previous super-resolution studies. Additionally, some Pom121 structures are resolved at the NE consistent with the composition of normal NPC’s, but with fewer localizations of Nup96 and Nup62 (**Fig. 2C-D**, circles). Considering that Pom121 acts as a deposition site of the NPC, these structures might correspond to the NPC’s observed at the early stage of their deposition to the NE, providing first insights into the NPC assembly pathway. Regarding lamin B1, which is a type V intermediate filament in the nuclear lamina that lines the inner surface of the NE relevant for chromatin organization^47^, previous SMLM studies showed that it is close to the nuclear membrane but its localization with respect to the NPC’s remained unclear^48^. Here we show with precise three-color imaging that it localizes very closely to the nuclear membrane and next to the NPC component Nup96 as visible in the “side views” (**Fig. 4B**).

Taken together, the analysis of NPC components Nup96, Nup62, Pom121 and lamin B1 illustrates the strength of our spectral demixing method for high-precision multi-color localization microscopy of challenging, multi-component cellular complexes. The splitSMLM method will be widely applicable, e.g. for a multitude of colocalization studies requiring multi-color SMLM and super-resolution imaging in biology. The SplitViSu software is made available under https://github.com/andronovl/SharpViSu.

## Methods

### Cell culture and immunofluorescence

U-2 OS-CRISPR-NUP96-mEGFP cells^34^ (300174, CLS) were plated in glass-bottom petri dishes (Cellvis D35-14-1.5-N). At ∼50% confluency the cells were washed twice in phosphate-buffered saline solution (PBS), fixed with 1% formaldehyde in PBS for 15 min and washed three times in PBS. The cells were permeabilized with 0.1% Triton X-100 in PBS (PBS/Tx) for 15 min and then blocked with 3% bovine serum albumin (BSA) in PBS/Tx (PBBx) for 1 hour. The primary antibodies were incubated, unless otherwise stated, at 2 µg/ml in PBBx overnight at +4 °C. The samples were washed three times with PBBx for at least 5 min each time. Then the secondary antibodies were incubated at 4 µg/ml in PBBx for 2 hours and the cells were washed again three times with PBS/Tx for at least 5 min each time. The samples were then postfixed with 1% formaldehyde in PBS for 10 min, washed twice with PBS and kept in PBS at +4 °C until mounting for imaging.

The following antibodies were used: Mouse anti-Nup62 (610497, BD Biosciences); Rabbit anti-GFP (TP-401, Torrey Pines Biolabs); Chicken anti-GFP (GFP-1010, Aves Labs); Mouse anti-GFP (2A3, IGBMC); Rabbit anti-lamin B1 (12987-1-AP, Proteintech), used at 0.7 µg/ml; Rabbit anti-Pom121 (GTX102128, GeneTex); Goat anti-Mouse CF680-conjugated (SAB4600199, Sigma); Goat anti-Rabbit CF680-conjugated (SAB4600200, Sigma); Goat anti-Rabbit CF660C-conjugated (SAB4600454, Sigma); Goat anti-Mouse CF660C-conjugated (SAB4600451, Sigma); Goat anti-Chicken CF660C-conjugated (SAB4600458, Sigma); Goat anti-Rabbit Alexa Fluor 647-conjugated (A-21245, Thermo Fisher); Goat anti-Mouse Alexa Fluor 647-conjugated (A-21236, Thermo Fisher).

### Mounting media

The samples were imaged in a water-based buffer that contained 200 U/ml glucose oxidase, 1000 U/ml catalase, 10% w/v glucose, 200 mM Tris-HCl pH 8.0, 10 mM NaCl, 50 mM MEA and with or without 2 mM COT. The mixture of 4 kU/ml glucose oxidase (G2133, Sigma) and 20 kU/ml catalase (C1345, Sigma) was stored at −20 °C in an aqueous buffer containing 25 mM KCl, 4 mM TCEP, 50% v/v glycerol and 22 mM Tris-HCl pH 7.0. MEA-HCl (30080, Sigma) was stored at a concentration of 1M in H_2_O at −20 °C. COT (138924, Sigma) was stored at 200 mM in dimethyl sulfoxide at −20 °C.

The samples were mounted immediately prior imaging using ∼200 µl of the imaging buffer and placing a clean coverslip on top of it while avoiding bubbles. The interface between the bottom of the petri dish and the clean coverslip in a well with imaging buffer is sufficiently airtight thanks to the surface tension and allows for long imaging runs (≥ 8 hours) without pH shift. After imaging, the samples were washed once with PBS and kept in PBS at +4 °C.

### Super-resolution imaging

The SMLM imaging was performed on a modified Leica SR GSD system (**Suppl. Fig. S1**). We used the HCX PL APO 160x/1.43 Oil CORR TIRF PIFOC objective that provides an equivalent pixel size of 100 nm on the camera. The fluorescence was excited with 642 nm 500 mW fiber laser (MBP Communication Inc.) and reactivated with 405 nm 50 mW diode laser (Coherent Inc.). We used a custom double-band filter cube (ordered and assembled at AHF Analysentechnik AG) with a dichroic mirror (DM1) Semrock FF545/650-Di01 and emission filters (F1) Semrock BLP01-532R mounted (airspaced) together with Chroma ZET635NF. The fluorescence was split into two channels with the Optosplit II (Cairn Research) image splitter, attached to a camera port of the Leica DMI6000B microscope. After splitting with the Chroma T690LPXXR dichroic mirror (DM2) and filtering the λ_S_ channel with the Chroma ET685/70m bandpass filter (F2), both channels were projected side-by-side onto the Andor iXon Ultra 897 (DU-897U-CS0-#BV) EMCCD camera. For measurements without image splitter, after the filter cube, the fluorescence was additionally filtered with the Semrock BLP01-635R-25 long-pass filter (F3) and the images were recorded with the Andor iXon+ (DU-897D-C00-#BV) EMCCD camera, attached to the second camera port of the microscope.

For SMLM imaging, the sample was illuminated with a constant power (30-50%) of the 642 nm laser. The first seconds were not recorded to avoid the initial high density of localizations. The acquisitions were started manually after appearance of separate single-fluorophore events (“blinking”). The exposure time of the camera was 8-13 ms; the electron multiplying gain of the camera was 300x (iXon Ultra) or 100x (iXon+). After 10-20 minutes, as the number of localizations dropped, the sample started to be illuminated additionally with a 405 nm laser with gradual increase of its intensity in order to keep a nearly constant rate of single-molecular returns into the ground state. The acquisition was stopped after almost complete bleaching of the fluorophore. Slight axial drifts were compensated by manual re-focusing, keeping in focus the localizations originating from the nuclear envelope.

For comparison between the imaging buffers, exactly the same parameters were kept for all acquisitions: the Andor iXon+ camera, EM gain 100x, exposure time 12.117 ms, 642 nm laser at 40% power. The first 100000 frames were acquired only with the 642 nm laser excitation. For the reactivation experiments, the same region was acquired a second time after the first acquisition. The intensity of the 405 nm laser was set to 0% at the beginning and then increased to 4% at about frame #1000, to 4.5% at frame #13000, 5% at #20000, 6% at #30000, 7% at #40000, 8% at #50000, 10% at #60000, 15% at #70000, 20% at #75000, 30% at #80000, 50% at #85000, 75% at #90000, 100% at #95000. The power of the 405 nm laser in % was set in the LAS X wizard, it does not correspond linearly to the real laser power, but provides the same power at same values, that is sufficient for comparison between different mounting media.

### Data processing

The localization and fitting of single-molecule events were performed in the Leica LAS X software with the “direct fit” fitting method, using an appropriate detection threshold. The localization tables were then exported for further processing in the SharpViSu software workflow^33^ and with customized Matlab procedures as implemented in SplitViSu. If imaged with the image splitter, the localizations were first demixed with the SplitViSu plugin and then processed in the main SharpViSu software for drift correction and relocalization on consecutive frames. Since the drift is the same for all unmixed channels, for drift correction in SharpViSu, we added possibilities to calculate the drift based on data in one of the channels or on the sum of data in all channels. The calculated by cross-correlation drift is then subtracted from localizations of each channel. For the data in this paper, we calculated the drift based on the sum of localizations in all channels.

For comparison between the imaging buffers (**Suppl. Fig. S8-S9**), all localizations were detected using a threshold of 25 photons/pixel in the LAS X wizard. After drift correction, the datasets were cropped to include only the surface of a nucleus with labelled nuclear pores (one nucleus per dataset). The photon counts and other localization statistics were calculated within these cropped regions, and the occurrence of localizations were normalized on the area of the corresponding regions, in order to get densities comparable between various datasets. All detected localizations within the region were assessed, without correction for relocalizations in consecutive frames and without any filtering. For all other datasets, we used a search radius of 50 nm for finding the localizations in the consecutive frames with their subsequent refinement. The resulting super-resolution images were built as 2D histograms of the single-molecule coordinates.

The emission spectra of fluorophores, observed in the λ_S_ or the λ_L_ channels, were calculated as a multiplication of the normalized emission spectrum of the fluorophores with the transmission of the dichroic mirrors and filters used in the given channel and the quantum yield of the camera. The average emission wavelength in a given spectral channel was calculated as 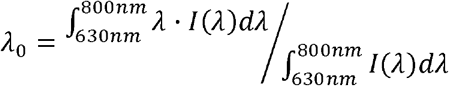, where λ is the wavelength and I(λ) is the emission of a fluorophore, observed in the given channel. For the case without image splitter, I(λ) was calculated considering the emission spectra of the fluorophores, transmission spectra of DM1 & F1 and the quantum yield of the camera.

FRC resolution (**Fig. 2E**) was calculated in SharpViSu^33^ according to the FRC_1/7th_ criterion^36^, using the histogram image reconstruction method with a pixel size of 5 nm and 90 frequency steps. The calculation was repeated 10 times for each dataset and the resolution was represented as a mean ± standard deviation of the obtained values for the corresponding image.

## Author contribution statement

L.A. designed research, software and analyzed data. L.A. and D.H. set up the optics. R.G. and L.A. performed fluorescence labelling and imaging. L.A. and B.P.K. wrote the manuscript with input from all authors. B.P.K. supervised the study.

## Additional Information

### Competing financial interests

The authors declare no competing financial interests.

## Acknowledgements

We thank Yongrong Liao and Izabela Sumara for anti-Pom121 antibodies; Amelie Zachayus and Arnaud Poterszman for the U2OS-NUP96-mEGFP cell line; Yvez Lutz (IGBMC Imaging Platform) for optimization of the labelling protocols & training. This work was supported by the French Infrastructure for Integrated Structural Biology (FRISBI) ANR-10-INSB-05-01, the Alsace Region, the infrastructures Instruct-ERIC and iNEXT-Discovery, the CNRS, Association pour la Recherche sur le Cancer (ARC), Institut National du Cancer (INCa), the Fondation pour la Recherche Médicale (FRM), and USIAS of the University of Strasbourg (USIAS-2018-012).

## Notes

### Competing Interest Statement

The authors have declared no competing interest.

