## Supplementary Figures for "A spectral demixing method for high-precision multi-color localization microscopy"

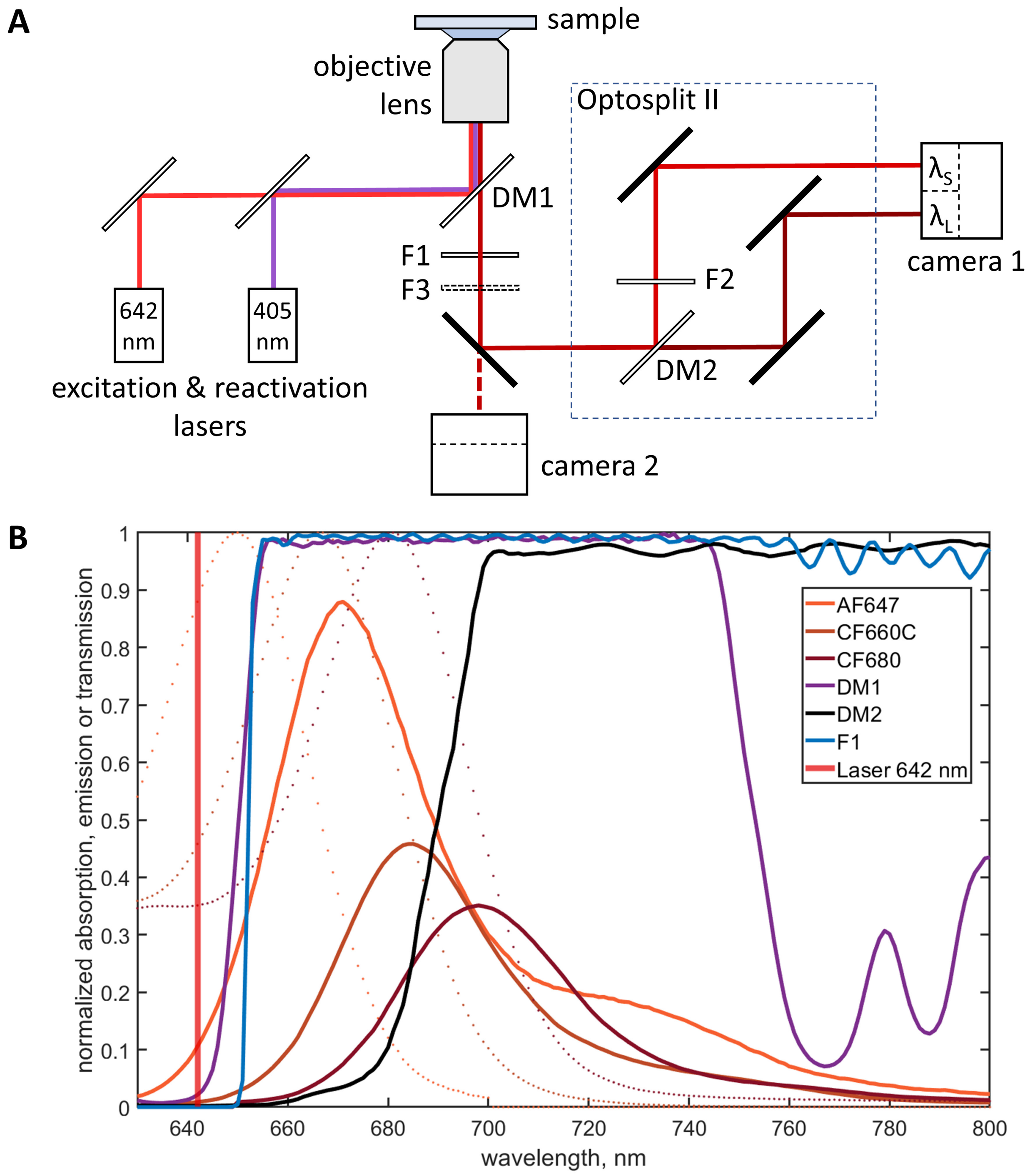


**Suppl. Fig. S1 Optical scheme and spectra**

(**A**) Optical scheme of the splitSMLM system. (**B**) Spectra of fluorophores and filters used in this study. The emission spectra of the fluorophores are normalized with respect to their excitation laser wavelenght.

| | AF647 | spectral difference ↔ | CF660C | spectral difference ↔ | CF680 | spectral difference ↔ | AF647 |
|---|---|---|---|---|---|---|---|
| Average emission wavelength | 686.31 nm | | 692.60 nm | | 702.82 nm | | |
| No image splitter | | 6.29 nm | | 10.22 nm | | 16.51 nm | |
| Average emission in $\lambda_S$ | 671.78 nm | | 677.95 nm | | 682.93 nm | | |
| All from $\lambda_S$ | | 6.17 nm | | 4.98 nm | | 11.16 nm | |
| Average emission in $\lambda_L$ | 711.63 nm | | 707.28 nm | | 710.36 nm | | |
| All from $\lambda_L$ | | 4.36 nm | | 3.09 nm | | 1.27 nm | |
| From brightest channel | | 35.50 nm | | 32.41 nm | | 38.58 nm | |
| Weighted mean | | 6.27 nm | | 10.17 nm | | 16.44 nm | |
| Simple mean | | 0.91 nm | | 4.03 nm | | 4.94 nm | |
| Theoretical $r = I_L / I_S$ | 0.57 | | 0.99 | | 2.59 | | |
| Experimental $r = I_L / I_S$ | 0.48 | | 1.11 | | 3.23 | | |

**Suppl. Fig. S2 Spectral characteristics of fluorophores with different demixing methods**

The blue line indicates similar spectral differences between the fluorophores when using the "weighted mean" method and without image splitter.

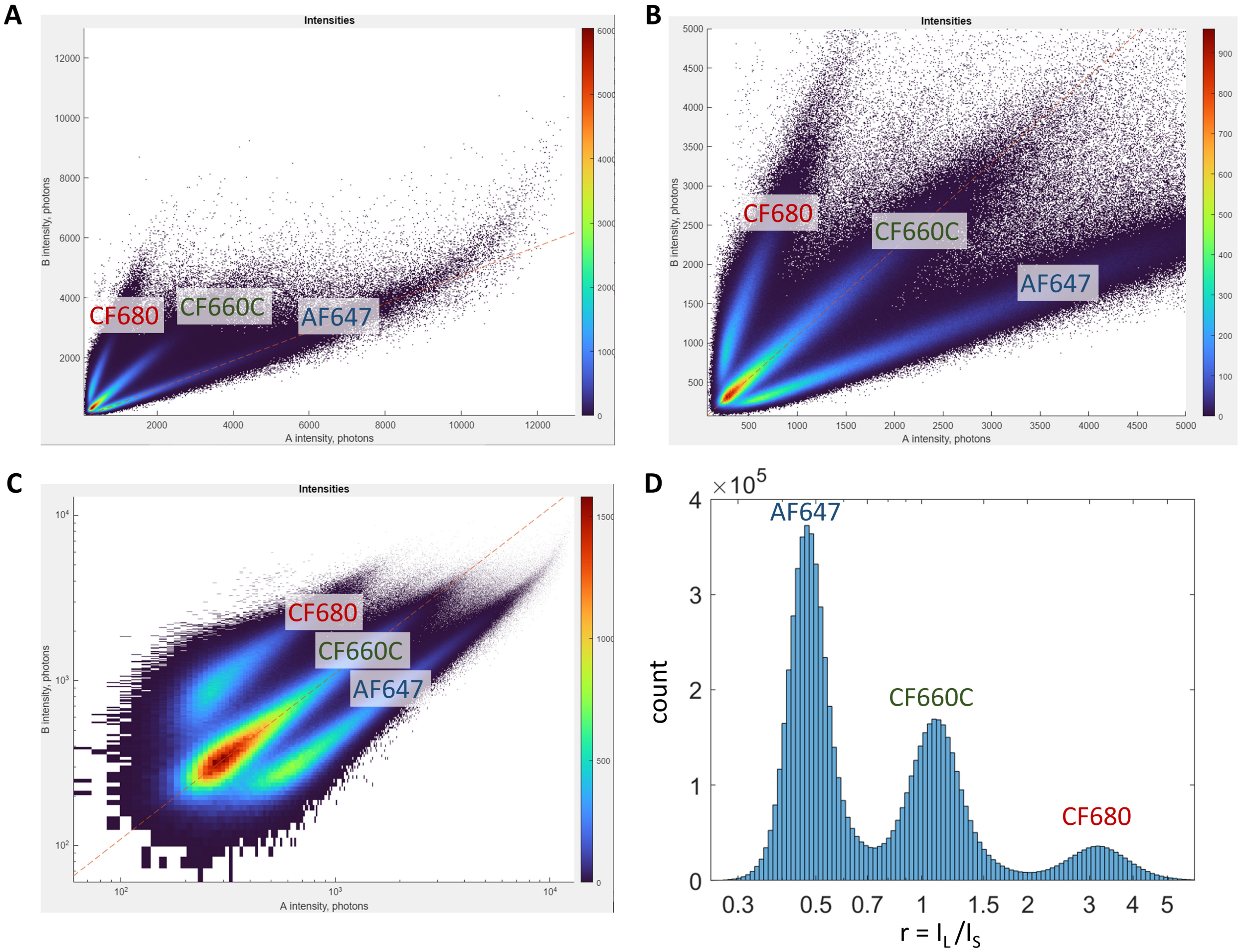


**Suppl. Fig. S3 Representation of splitSMLM data**

(**A-C**) Bivariate histograms of intensities in the $\lambda_L$ and $\lambda_S$ channels, showing the full extent of the intensities (A), only the localizations with less than 5000 photons (B) and full extent of the data displayed as a log-log plot (C). (**D**) Univariate histogram of ratios *r* on a semi-log plot with peaks corresponding to the different fluorophore species.

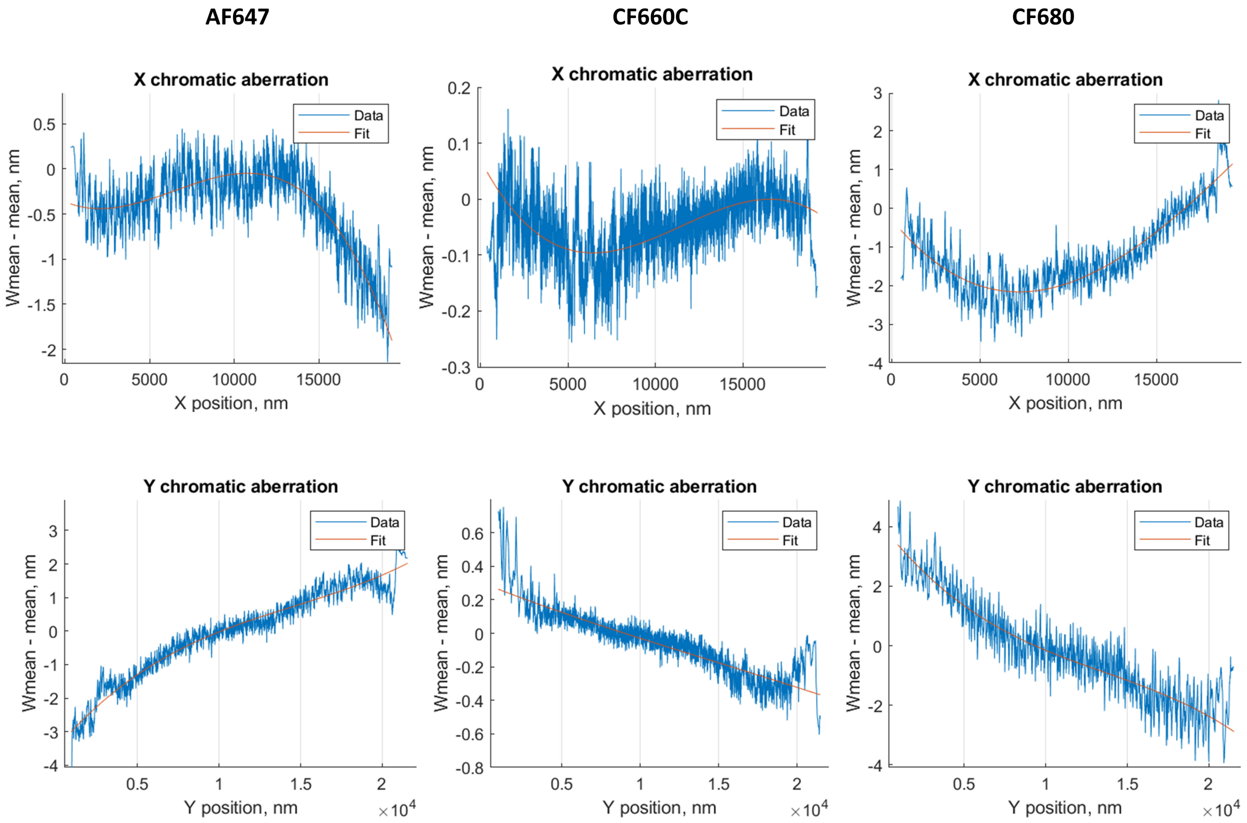


**Suppl. Fig. S4 Correction of chromatic aberrations in SplitViSu**

The graphs show residual chromatic shift between the coordinates of the fluorophores, calculated as the “weighted mean” and the “simple mean” of the input coordinates (blue, using a moving average window to reduce fluctuations) and its polynomial fit (red curves).

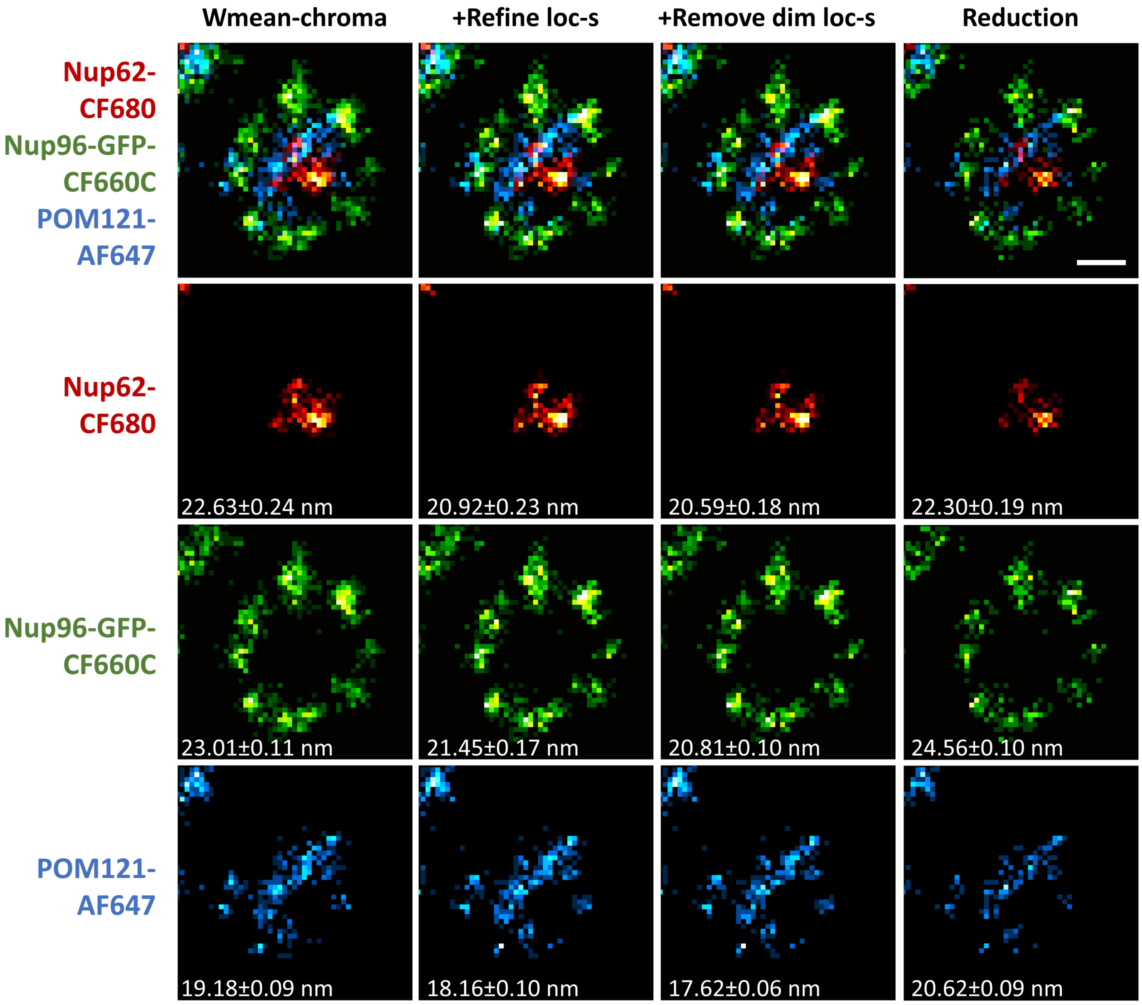


**Suppl. Fig. S5 Refinement of multiple localizations**

Color coding of the labeled proteins as in Fig. 2. $FRC_{1/7th}$ resolution values are shown at the bottom. Scale bar, 50 nm.

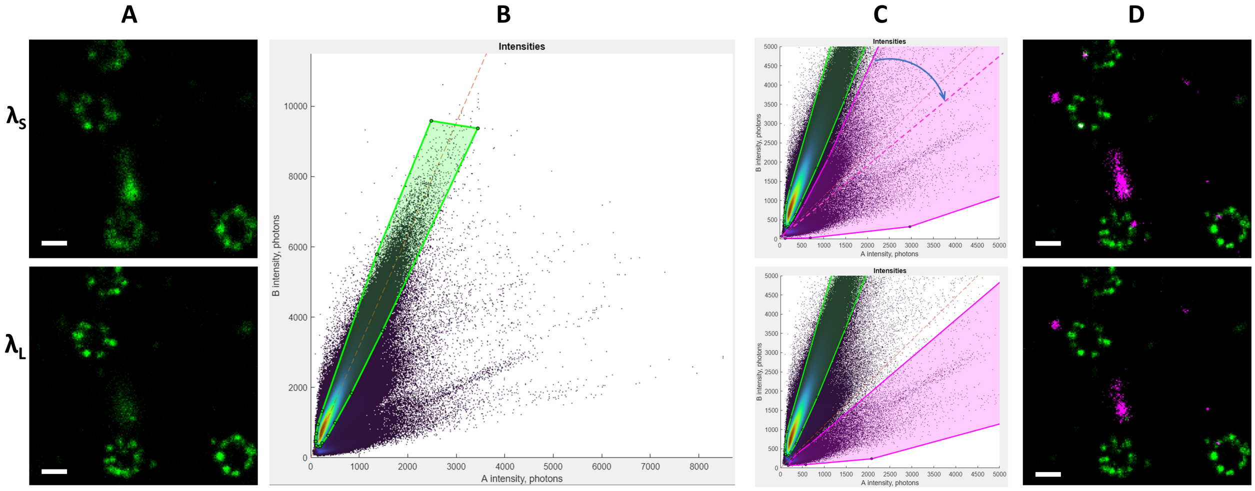


**Suppl. Fig. S6 Removal of spurious localizations with splitSMLM**

(**A**) SR images of a single-labelled sample Nup96-AF647, reconstructed from localizations in the $\lambda_S$ or $\lambda_L$ channels. (**B**) Bivariate histogram of photon counts of this sample with the green region corresponding to AF647. (**C**) Spectral demixing using a region for AF647 (green) and two different regions for spurious localizations (magenta). (**D**) Demixing in SplitViSu using the corresponding regions in (C) allows separation of reliable signal (green) from spurious localizations (magenta).

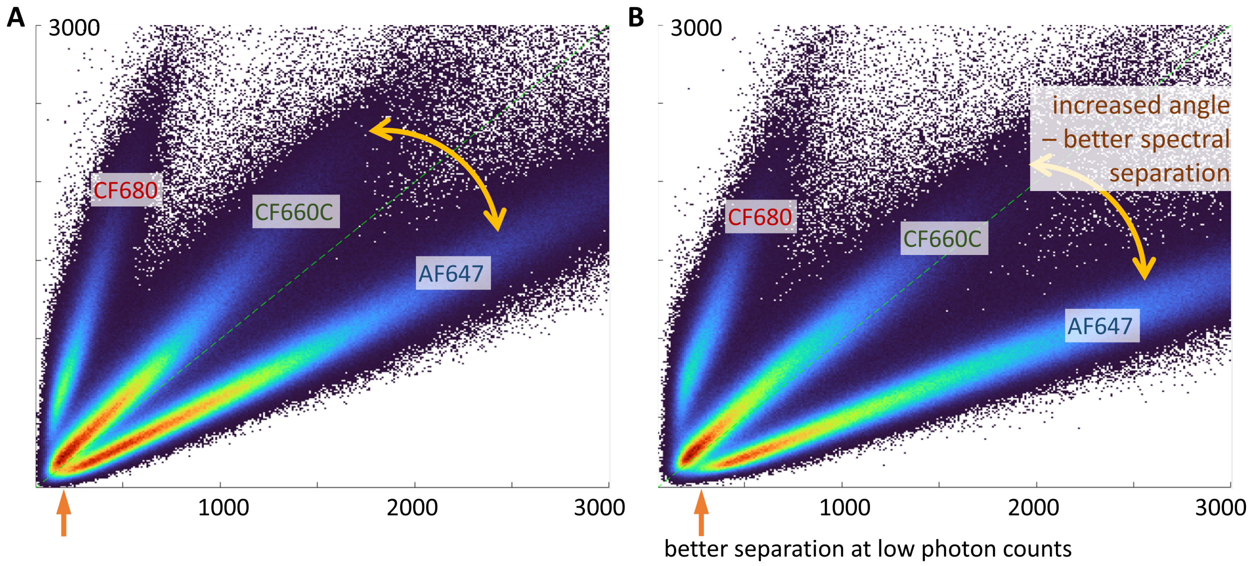


**Suppl. Fig. S7 Improvement of the spectral separation of fluorophores with dichroic filters**

(**A**) Bivariate histogram of photon counts of a triple-labelled sample using a standard filter cube Leica GSD 642HP-T, consisting of an excitation filter zet405_642x, a dichroic mirror ZT405/642rpc and an emission filters – combination of et710_100lp and ET650LP. (**B**) Bivariate histogram of photon counts of the same sample using our custom filter cube allows better separation of dim fluorophores due to detection of fluorescence close to the laser line.

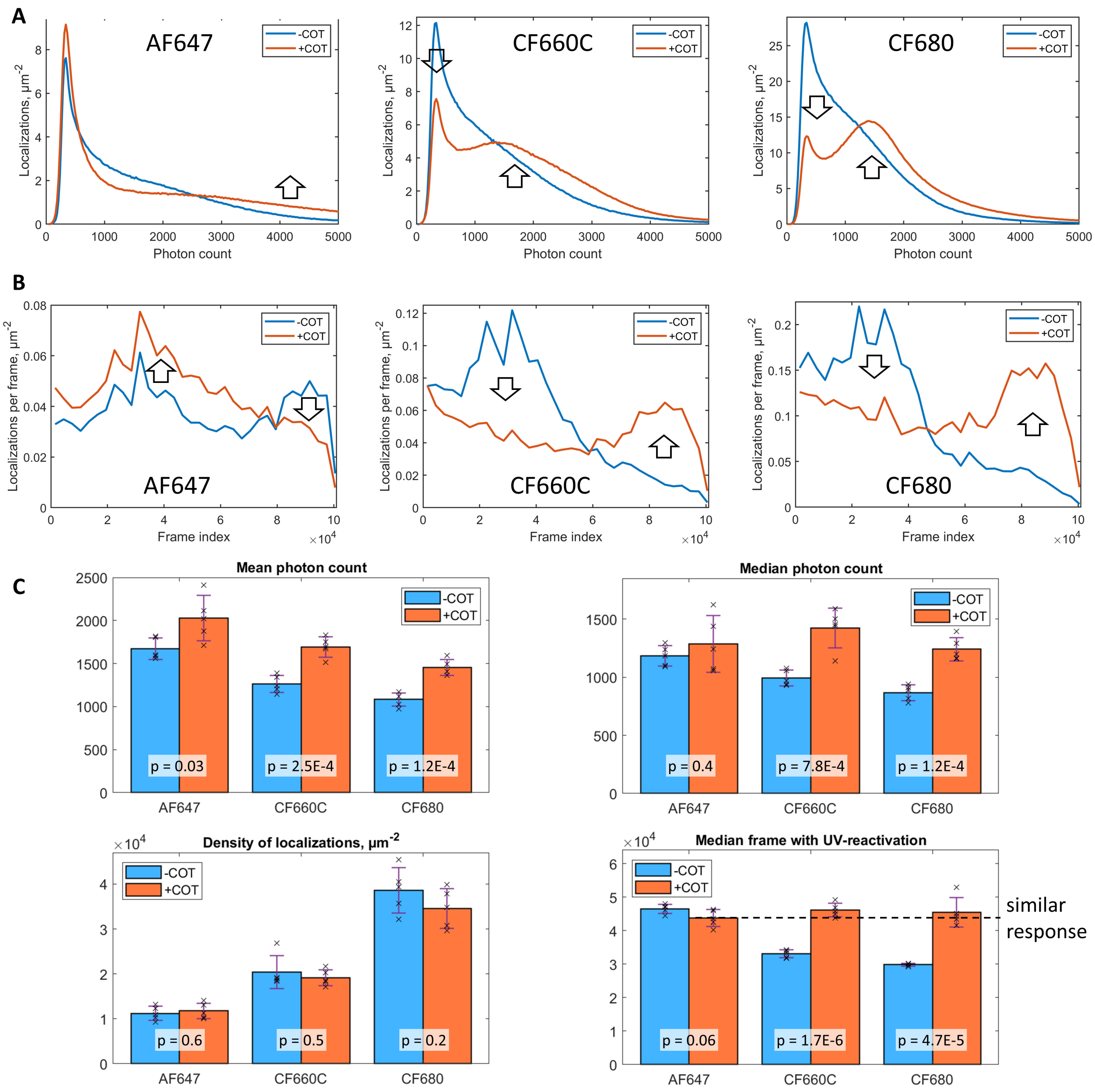


**Suppl. Fig. S8 Behavior of fluorophores in a COT-supplemented imaging buffer**

(**A**) Typical photon counts of fluorophores in a buffer with (red) or without (blue) COT. (**B**) Response on reactivation light – number of localizations per frame with stepped increase of the 405 nm laser power in a buffer with (red) or without (blue) COT. (**C**) Statistics of fluorophore behavior, demonstrating a significant increase in the photon count in a buffer with COT for all three fluorophores, while preserving the density of localizations and therefore equalizing the fluorophore response on reactivation with 405 nm light.

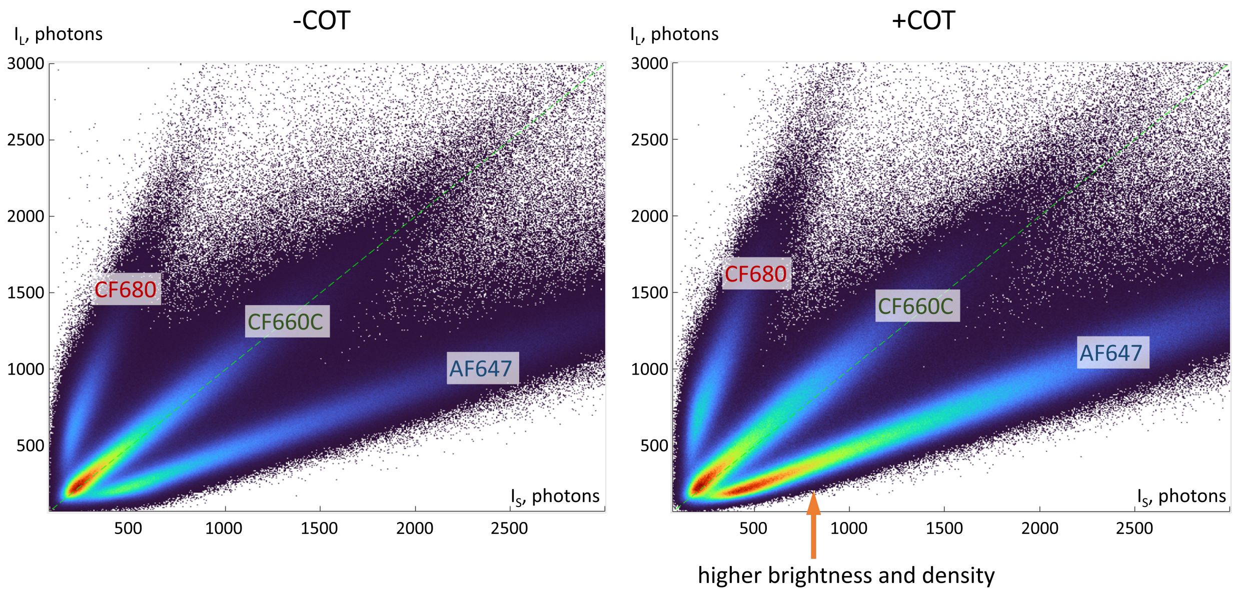


**Suppl. Fig. S9 Bivariate histogram of photon counts**

Bivariate histogram of photon counts of a triple-labelled sample in an imaging buffer without COT (left) or supplemented with 2 mM COT (right).

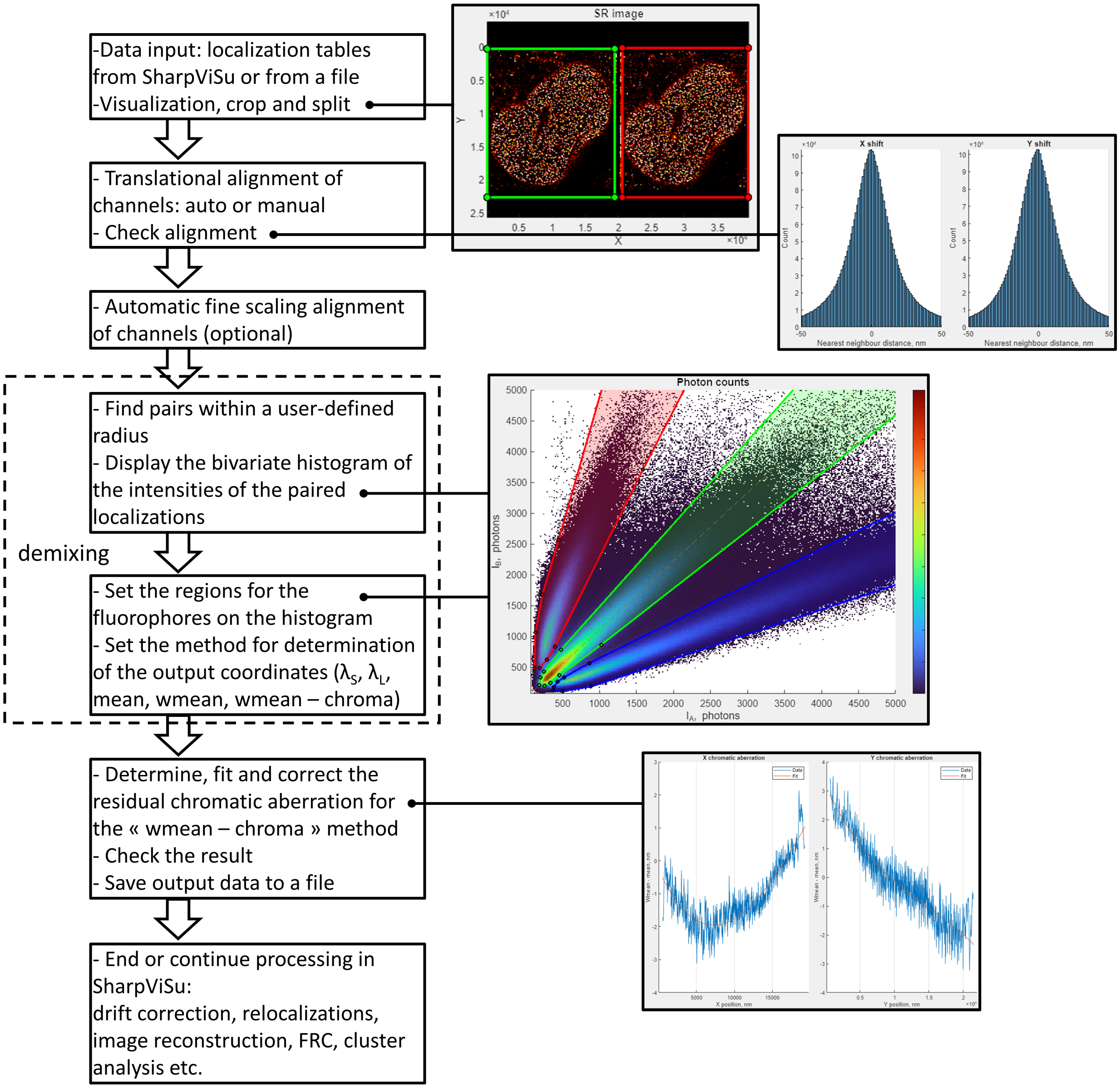


**Suppl. Fig. S10 Workflow of fluorophore demixing in SplitViSu**
